# Weak, specific chemical interactions dictate barnase stability in diverse cellular environments

**DOI:** 10.1101/2024.09.26.615174

**Authors:** Ume Tahir, Caitlin M. Davis

## Abstract

It is well-established that in vitro measurements do not reflect protein behaviors in-cell, where macromolecular crowding and chemical interactions modulate protein stability and kinetics. Recent work suggests that peptides and small proteins experience the cellular environment differently from larger proteins, as their small sizes leave them primarily susceptible to chemical interactions. Here, we investigate this principle in diverse cellular environments, different intracellular compartments and host organisms. Our small protein folding model is barnase, a bacterial ribonuclease that has been extensively characterized in vitro. Using fast relaxation imaging, we find that FRET-labeled barnase is stabilized in the cytoplasm and destabilized in the nucleus of U2-OS cells. These trends could not be reproduced in vitro by Ficoll and M-PER^TM^, which mimic macromolecular crowding and non-specific chemical interactions, respectively. Instead, in-cell trends were best replicated by cytoplasmic and nuclear lysates, indicating that weak specific interactions with proteins in either compartment are responsible for the in-cell observations. Interestingly, in the cytoplasm barnase’s unfolded state is unstable and prone to aggregation, while in the nucleus a stable unfolded state exists prior to aggregation. In the more biologically relevant environment of bacterial cells, barnase folding resembled that in the nucleus, but with no aggregation at higher temperatures. These findings show that protein interactions are evolved for their native environment, which highlights the importance of studying and designing proteins in situ.

## 1 Introduction

It is well-established that macromolecular crowding and chemical interactions between biomolecules inside cells can substantially tune both protein stability and folding kinetics (Danielsson et al., 2015; Davis & Gruebele, 2018; Ebbinghaus et al., 2010; Knab & Davis, 2023; Monteith et al., 2015). Furthermore, given the heterogeneity of the cell interior, these properties may vary depending on the local microenvironment. For example, it has been demonstrated that two model proteins, phosphoglycerate kinase (PGK) and variable major protein-like sequence expressed (VlsE), exhibit different stabilities and folding kinetics in different cellular compartments (Dhar et al., 2011; Tai et al., 2016). Thus, in-cell measurements contain information in a functionally-relevant context that cannot be simply extrapolated from in vitro data.

Currently, much of our understanding of how the cellular environment modulates protein behavior, especially in the context of eukaryotic cells, is derived from large proteins, like PGK and VlsE. There is comparatively less known about how peptides and small proteins behave in eukaryotes, despite their diverse biological functions – including as enzymes, chaperones, and regulators of cell division (Steinberg & Koch, 2021; Storz et al., 2014). Given their small sizes relative to the surrounding macromolecules, peptides and small proteins are expected to experience the cellular environment differently from larger proteins. While the average ≍400 amino acid eukaryotic protein is susceptible to both macromolecular crowding and chemical interactions, peptides and small proteins are unlikely to be sterically constrained by larger macromolecules (Brocchieri & Karlin, 2005; Christiansen et al., 2010; Chu et al., 2023). Indeed, recent work on the 80 amino acid λ-repressor protein fragment (λ_6-85_), suggests that the stability of small proteins in eukaryotic cells is dictated solely by chemical interactions (Knab & Davis, 2023). This aligns with earlier results for SH3, a small protein folding model that is destabilized by chemical interactions in both prokaryotic and eukaryotic cells, despite the vastly different concentrations of biomolecules in these environments (Smith et al., 2016; Thole et al., 2021). However, there is still a need for additional studies to determine if this principle applies to other small proteins and to further explore the effects of different types of cellular microenvironments on small protein and peptide stability.

This work focuses on barnase (Figure 1A), a 110-residue extracellular ribonuclease from *Bacillus amyloliquefaciens* (Hartley, 1989). Its fold and function are of interest both in prokaryotic cells where it natively exists, and eukaryotic cells where it may be a promising candidate for anticancer therapy (Shilova et al., 2021; Ulyanova et al., 2011). Barnase has an extensive history of being used as a protein folding model, with a thoroughly characterized in vitro folding mechanism (Alemany et al., 2016; Fersht, 1993, 2000; Kippen et al., 1994). It has contributed to our understanding of the principles that govern protein folding, with systematic mutations providing residue-level details on how noncovalent interactions drive higher-order structure formation, and generally aided in method development for elucidating protein folding pathways, including detection of intermediates (Fersht, 2000; Khan et al., 2003; Kippen et al., 1994; Serrano et al., 1992). Intracellularly, barnase is tightly bound to its inhibitor, barstar; this complex has similarly become a paradigm for studying protein-protein interactions (Hartley, 1993; Lee & Tidor, 2001; Schreiber et al., 1994). The abundance of in vitro data makes barnase an interesting candidate for investigating how the cellular environment tunes protein behavior. For example, a collection of barnase mutants with differing stabilities has been used as biosensors to report on in-cell chaperone activity (Wood et al., 2018). With Förster resonance energy transfer (FRET) reporting on the folding state of the protein, it was found that the barnase mutants have different distributions between the folded, unfolded, and aggregated states in the cytoplasm and nucleus (Raeburn et al., 2022). These differences in behavior across different cellular compartments align with the knowledge that barnase, both independently and as part of a complex with barstar, is sensitive to its local microenvironment; its stability is known to be variable with salt identity and concentration, and complex formation is facilitated by cellular crowding effects (Bye et al., 2016; Qi et al., 2014; Ridgway et al., 2008; Trevitt et al., 2024). These observations further raise the question of how the local microenvironment modulates the folding of small proteins and peptides, both in prokaryotic and eukaryotic cells.

**Figure 1.**
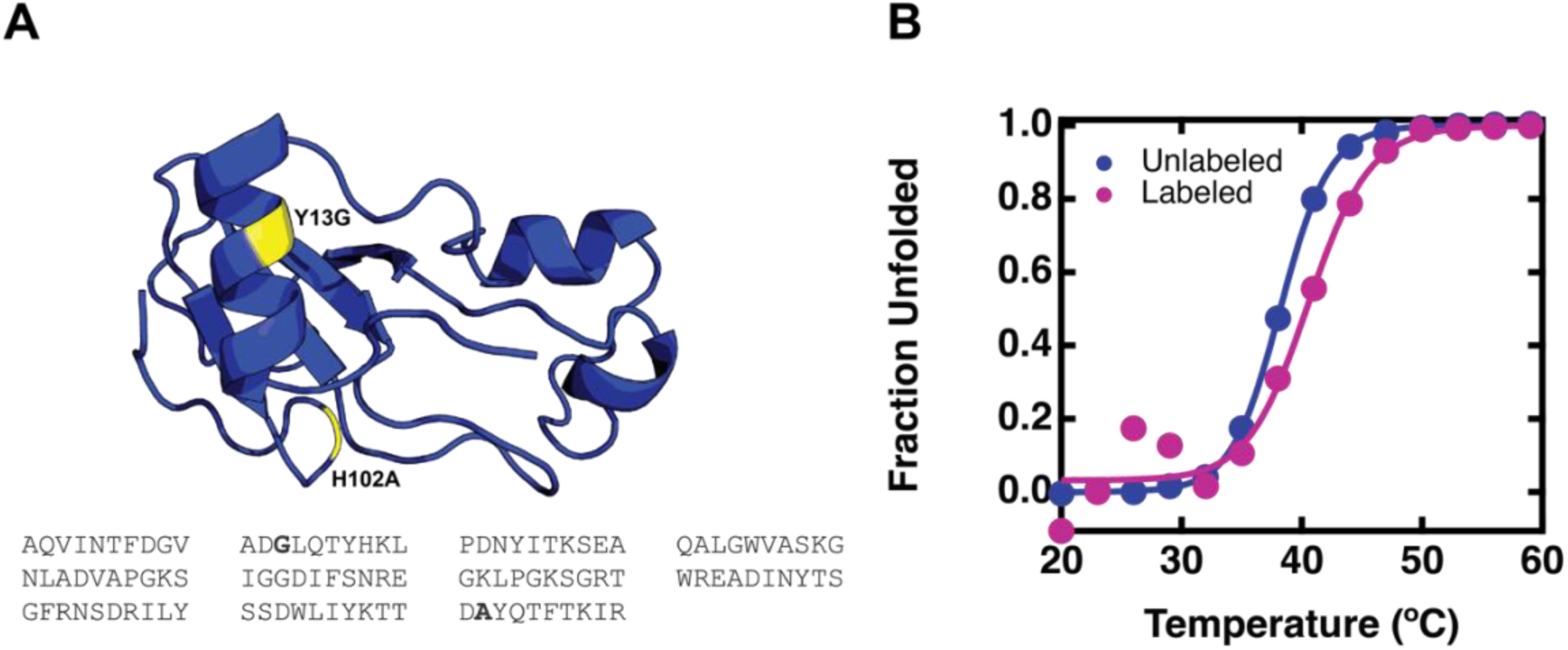
(A) Structure and sequence of barnase with Y13G and H102A mutations highlighted in yellow (PDB: 1BNR). Figure created using PyMOL (The PyMOL Molecular Graphics System, n.d.). (B) Representative fluorescence melts for 10 μM unlabeled (blue) and 0.25 μM FRET-labeled (magenta) barnase in 20 mM sodium phosphate, pH 7.4. The melts were normalized and fit to a two-state denaturation (E1).

Here, we focus on barnase as a small protein folding model and explore the effect of different cellular compartments on its stability. We find that compared to in vitro, barnase is slightly stabilized in the cytoplasm and significantly destabilized in the nucleus of mammalian U2-OS cells. Interestingly, a stable unfolded state exists in the nucleus, whereas barnase is more aggregation-prone in the cytoplasm. To understand the differences between in vitro and in-cell stabilities, we used Ficoll and mammalian protein extraction reagent (M-PER^TM^) to mimic crowding and non-specific chemical interactions, respectively, and found that neither could replicate cellular trends. Instead, these trends were best reproduced by cytoplasmic and nuclear lysates, suggesting that weak specific interactions with proteins in either compartment are most important for determining cellular stability. Lastly, we investigated how the conclusions drawn from eukaryotic U2-OS cells translate to the more biologically relevant context of bacterial cells. We found that unlike in mammalian cells, barnase does not aggregate upon unfolding in bacterial cells demonstrating that evolution has carefully optimized protein interactions in their native environment. This has important implications for functional protein design, which is typically performed in vitro, and interpretation of phylogenic reconstructions of ancestral proteins, which is often performed protein by protein.

## 2 Results and Discussion

### 2.1 The addition of FRET labels to barnase is minimally perturbative

For this work, we used the Y13G H102A mutant of barnase (Figure 1A). The H102A mutation is commonly used to diminish the toxic ribonuclease activity of barnase in the absence of its intracellular inhibitor (Meiering et al., 1992), and the Y13G mutation further destabilizes the construct to yield a melting temperature (T_M_) that is compatible with our in-cell temperature-dependent experiments (36.7 °C) (Vu et al., 2004). The effects of the cellular environment on protein folding can be described thermodynamically by the contributions of steric crowding and non-steric chemical interactions in the cell to the free energy of the folding reaction, Δ*G*° (Davis et al., 2018).

Steric crowding interactions are purely entropic, accounting for the volume that a protein occupies in solution (Zhou et al., 2008). Therefore, these interactions depend on the structure of the folded and unfolded state of a protein and not on its sequence. The active site H102A mutation is not structurally disruptive as confirmed by NMR (Vu et al., 2004). Additionally, the double mutant Y13G H102A has a circular dichroism spectrum that closely matches wildtype barnase (Vu et al., 2004; Vuilleumier et al., 1993). Therefore, the effects of steric crowding on the stability of barnase mutants should closely match the effects of steric crowding on wildtype barnase.

On the other hand, chemical interactions depend on the sequence of a protein, more specifically, the solvent accessible surface area of the folded and unfolded state. Chemical interactions are enthalpic and account for all other interactions a protein will have with its environment aside from steric crowding. A single point mutation can change the rotational motion of a protein by modulating its surface net charge density, surface hydrophilicity, and the electric dipole moment (Mu et al., 2017). However, because a single point mutation represents a small percentage of the overall surface of a protein, it is unlikely to overcome the chemical interactions of the rest of the protein (Feng et al., 2019; Monteith et al., 2015). Therefore, the effects of non-steric chemical interactions on the stability of barnase mutants will provide a qualitative trend for wildtype barnase.

Although the magnitude of stability changes observed in the cell, determined by the combination of steric and non-steric interactions, will differ between mutant and wildtype barnase sequences, the overall trends should be similar. To track the folding state of the barnase mutants in-cell, we designed a FRET-labeled construct with AcGFP1 as the donor fluorophore on the N-terminus and mCherry as the acceptor fluorophore on the C-terminus. The donor to acceptor fluorescence ratio (D/A) reports on the end-to-end distance of the protein, with higher values representing a more extended state. However, prior to using this construct in-cell, it was important to verify that the labels did not perturb the folding behavior of barnase. We expressed and purified the unlabeled construct and found its structure and T_M_ to be consistent with the literature using circular dichroism (Figure S1) (Vu et al., 2004; Vuilleumier et al., 1993). Next, we derived T_M_ values for the unlabeled and labeled constructs using tryptophan fluorescence and FRET melts, respectively. We found that the T_M_ of the unlabeled construct was 38.16 ± 0.07 °C, and the T_M_ of the FRET-labeled construct was 39 ± 2 °C (Figure 1B). We therefore concluded that the introduction of FRET labels was minimally thermodynamically perturbative.

### 2.2. Barnase is slightly stabilized in the cytoplasm and significantly destabilized in the nucleus of U2-OS cells

To examine the effect of different local cellular environments on barnase stability, we transfected U2-OS cells with FRET-labeled barnase containing either a nuclear export or nuclear localization sequence in U2-OS cells (Table S1). The stability in the cytoplasm and nucleus of U2-OS cells was probed by fast relaxation imaging (FReI), which couples the spatial resolution of fluorescence microscopy with perturbations in the form of laser-induced temperature jumps; thermodynamic information is derived from the 10 second equilibrium phase following the temperature jump (Ebbinghaus et al., 2010).

We observed distinct behaviors in the cytoplasm and nucleus. Interestingly, the D/A in the cytoplasm consistently trended downwards with increasing temperature, which is indicative of aggregation without an observable, stable unfolded state (Figure 2). Thus, we report an aggregation temperature (T_A_) in place of T_M_ for the cytoplasm. The T_A_ was 41.4 ± 0.5 °C across 70 cells (Table S2). This reflects that the folded state is slightly stabilized by ≍2 °C relative to in vitro.

**Figure 2.**
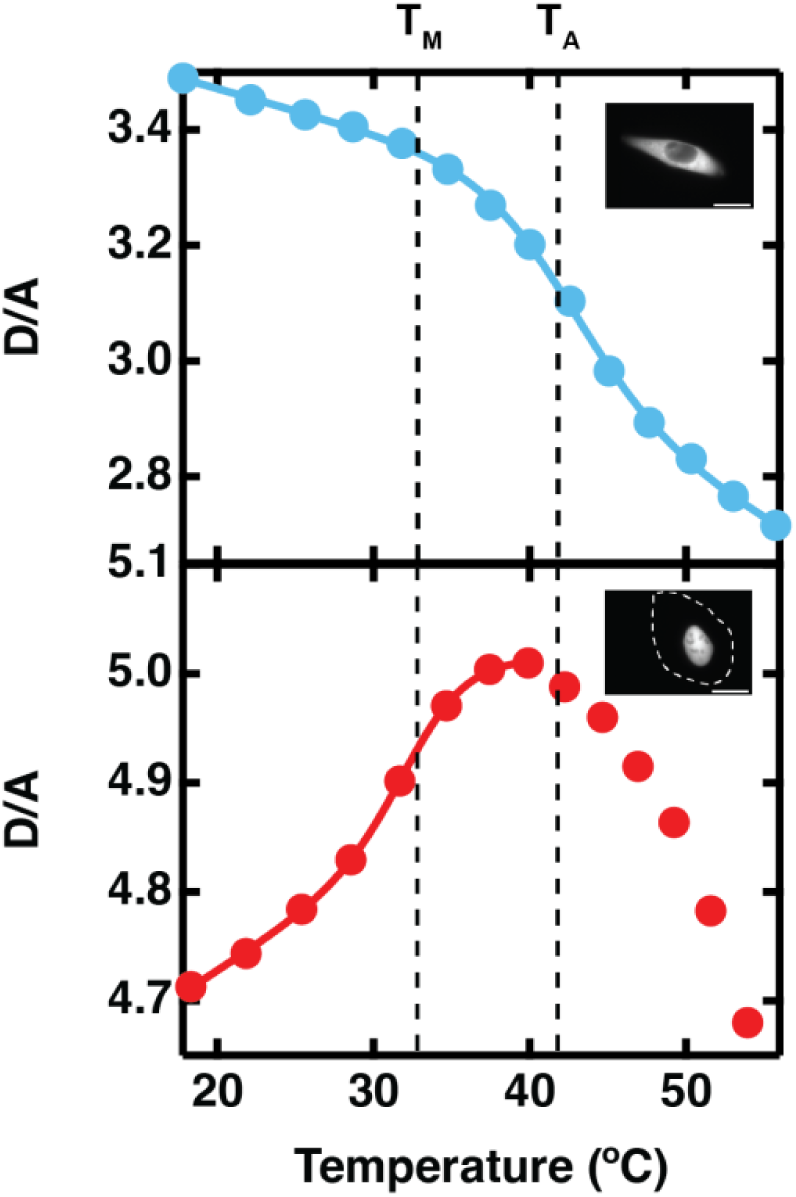
Representative donor/acceptor (D/A) melting curves extracted from FReI of barnase in the cytoplasm (top) and nucleus (bottom) of U2-OS cells. The data are fit to a two-state denaturation (E1) with dashed vertical lines indicating the melting temperature in the nucleus (T_M_) and midpoint of aggregation in the cytoplasm (T_A_), respectively. Insets display representative cell images. In the bottom image the cell membrane is highlighted by a white dashed line. Scale bar is set to 10 μm.

In the nucleus, there is a clear unfolding transition prior to the onset of aggregation at a T_A_ similar to the cytoplasm (Figure 2). The T_M_ was 26.8 ± 0.3 °C across 75 cells (Table S3). This reflects a significant destabilization of ≍12 °C relative to in vitro, and ≍15°C relative to the cytoplasm. A subset of cells (44%) exhibited only aggregation at T_A_; these cells tended to have a lower or greater than average D/A, suggesting that the initial conformation was either misfolded or unfolded (Figure S2). Barnase localized to the nucleus is predominantly unfolded but below T_A_ at normal cell culture temperatures (37°C), and therefore our FReI experiments that begin at room temperature (≍18 °C) are conducted on cells with populations of barnase that can be reversibly folded. In vitro, barnase undergoes reversible unfolding induced by heat and numerous other perturbants (Fersht, 1993; Johnson & Fersht, 1995). However, our FReI experiments demonstrate that interactions with the cellular environment of U2-OS cells decrease barnase’s thermal refoldability.

### 2.3. Cellular stability trends cannot be explained by macromolecular crowding or non-specific chemical interactions alone

The cell interior is primarily water, and the remaining 30-40% consists of ions, various small organics and metabolites, and macromolecules (Davis & Gruebele, 2018; Davis et al., 2018; Zimmerman & Trach, 1991). Therefore, protein stabilities in-cell may be modulated by factors like macromolecular crowding and chemical interactions. Macromolecular crowding entropically destabilizes the extended unfolded state of a protein, but may have minimal impact on smaller proteins, like barnase, that are not sterically confined by larger macromolecules (Knab & Davis, 2023). Additionally, the differing stabilities of barnase in the cytoplasm and nucleus of U2-OS cells are unlikely to be explained by macromolecular crowding theory, given the relatively conserved concentrations of macromolecules across these cellular compartments (Guigas et al., 2007). Instead, since the cytoplasm and nucleus vary in their chemical compositions it is more probable that stability differences arise from differences in chemical interactions. Chemical interactions account for the stabilizing, repulsive and destabilizing, attractive forces between molecules in the absence of entropic crowding (McConkey, 1982). Therefore, unlike macromolecular crowding, chemical interactions depend on the composition of the local environment as well as the physicochemical properties of the surface of a protein. Because the surface of a folded and unfolded protein differ, chemical interactions will affect the free energy of the folded and unfolded state differently and the net effect on the protein can be either stabilizing or destabilizing (Davis et al., 2018).

Cellular mimetic reagents can be used in vitro to untangle the origin of in-cell stability trends. Because of its relatively inert surface, Ficoll is often used to isolate crowding effects (Benton et al., 2012; Davis et al., 2020; Knab & Davis, 2023). Here, we derived melting temperatures for unlabeled barnase in the presence of increasing concentrations of Ficoll (Figure 3). We find that although there is a small stabilization trend between 0-35% Ficoll, which could account for the slight ≍2 °C stabilization of barnase in the cytoplasm, it does not explain the ≍12 °C destabilization in the similarly crowded nucleus. Additionally, the previously validated cytomimetic concentration of Ficoll (15%) (Davis et al., 2020; Knab & Davis, 2023), near the physiological concentration of macromolecules in eukaryotic cells, produces only a mild stabilization of ≍0.8 °C. Therefore, as predicted, it is unlikely that macromolecular crowding plays a large role in regulating barnase stability in-cell.

**Figure 3.**
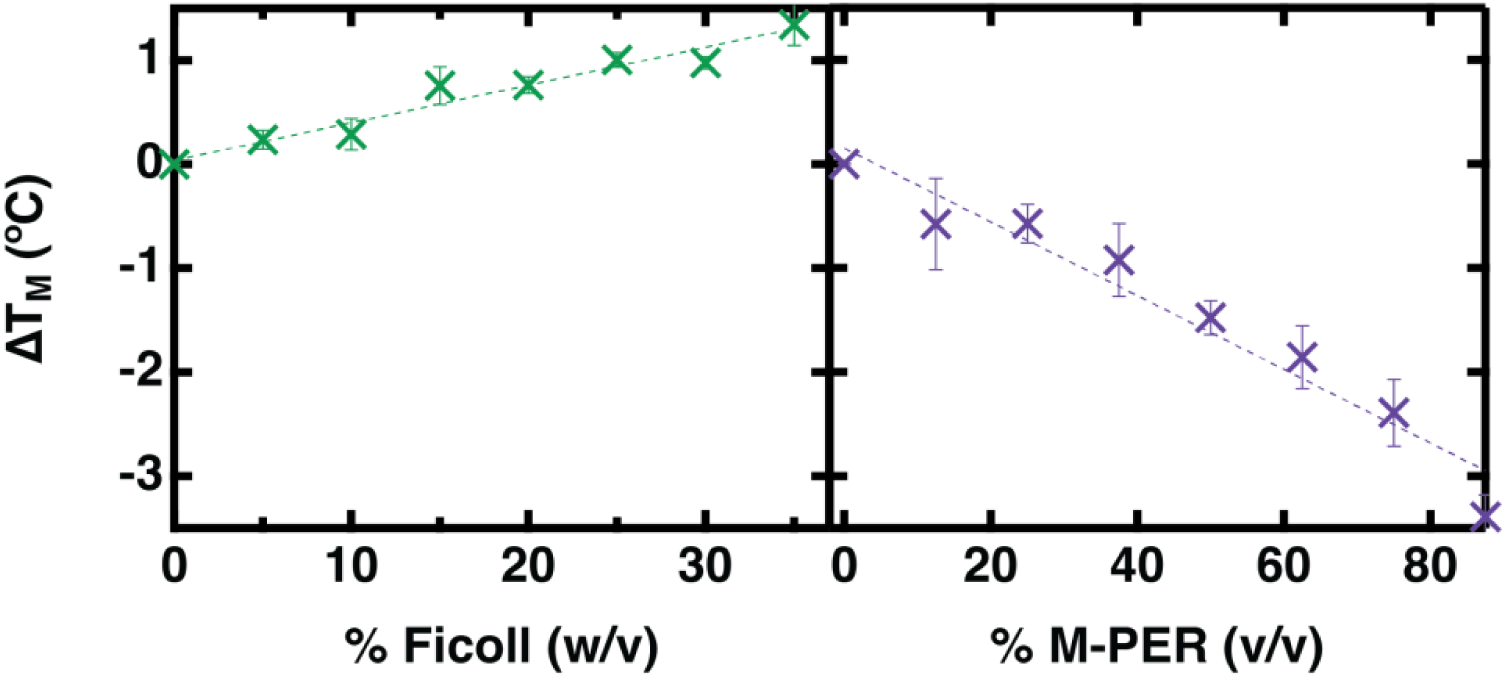
Change in the melting temperature of unlabeled barnase relative to 20 mM sodium phosphate, pH 7.4 with increasing concentrations of Ficoll and M-PER^TM^. Melting temperatures derived from tryptophan melts with 10 μM barnase. Error bars display standard deviation of 3 measurements. Data is fit to a line.

Other than crowding, proteins also experience non-specific chemical interactions in-cell with ions, small organic molecules, fatty acids, amino acids, etc. Ionic strength is important because the presence of ions may electrostatically screen a protein from interactions with surrounding molecules; here, we replicate intracellular ionic strength using 150 mM KCl, as K^+^ is the most abundant ion in mammalian cells (Ames & Nesbett, 1966). We find that although K^+^ slightly stabilizes barnase by 1.66 ± 0.06 °C, it does not fully recapitulate our in-cell observations. The concentration of K^+^ is likely higher in the nucleus than in the cytoplasm (Kellermayer, 1981; Paine et al., 1981), so this stabilizing effect does not account for the significant destabilization observed in the nucleus relative to the cytoplasm. Furthermore, prior work examining the Hofmeister effect on barnase stability found no significant changes under physiological salt conditions (Bye et al., 2016). In the spirit of *E. coli* work that reproduced the ionic and metabolic environment using the top 80% of cellular salts (Sieg et al., 2022), Eco80, we tested the stability of barnase in the top 80% metabolites identified in the cytoplasm of the HeLa cell line (Chen et al., 2016), HeLa80 (metabolic data is not available for U2-OS cells). We find that neither reproduces the observed cytosolic trends (Table S4).

We use M-PER^TM^ to account for non-specific chemical interactions with small molecules in the cell. M-PER^TM^ is a mild cell lysis reagent formulated to minimally interfere with downstream biological applications by preserving native protein properties. Specifically, M-PER^TM^ contains a proprietary zwitterionic detergent in 25 mM bicine buffer (pH 7.6) that mimics interactions with lipids and amino acids, respectively (Knab & Davis, 2023; Yoo & Davis, 2022). Converse from Ficoll, we find that there is a destabilization trend with increasing concentrations of M-PER^TM^ (Figure 3). However, at the previously validated cytomimetic concentration of 20% M-PER^TM^ (Knab & Davis, 2023; Yoo & Davis, 2022), the destabilization is only 0.5 °C compared to a ≍12 °C destabilization in the nucleus. Thus, while chemical interactions with small molecules may account for some of the nuclear destabilization of barnase, the observed cellular trends cannot be fully reproduced by either macromolecular crowding or non-specific chemical interactions alone.

### 2.4. Differences in stability between the cytoplasm and nucleus arise from weak specific interactions with proteins in either compartment

Although Ficoll and M-PER^TM^ can reproduce non-specific crowding and chemical interaction effects, they do not capture behaviors that arise from weak, transient but specific interactions with macromolecules in the local microenvironment in-cell. The cytoplasm and nucleus differ in the types and quantities of macromolecules present, with the cytoplasm containing machinery for diverse metabolic functions, including protein synthesis (Kırlı et al., 2015), and the nucleus having more specialized proteins, including highly charged histones and transcriptional machinery, in addition to nucleic acids (van der Zanden et al., 2023). Thus, to account for weak, specific chemical interactions in the cytosol we mimic the cytoplasmic environment in vitro using cytoplasmic lysate, which contains cytoplasmic proteins extracted from U2-OS cells; likewise, we mimic the specific-protein and non-specific nucleic acid components of the nucleus using nuclear lysate, which contains nuclear proteins extracted from U2-OS cells, and salmon sperm DNA, respectively.

We find that FRET-labeled barnase experiences a slight destabilization of 2.5 ± 0.5 °C in the presence of 1 mg/ml salmon sperm DNA, concentrations that are insufficient to induce macromolecular crowding (Figure 4, Table S5). Barnase is a ribonuclease that catalyzes the cleavage of single-stranded RNA. Positively charged residues at the binding interface of barnase bind with the negatively charged RNA substrate (Spaar et al., 2006). This positively charged patch is likely also poised to interact with other negatively charged biomolecules like DNA. We observe that barnase is destabilized by these non-specific, attractive interactions with DNA. This suggests that chemical interactions with nucleic acids may be contributing to the destabilization of barnase in the nucleus of U2-OS cells. However, it is likely that nonspecific interactions with RNA in the cytoplasm are also destabilizing. Thus, interactions with nucleic acids alone cannot recapitulate our in-cell observations.

**Figure 4.**
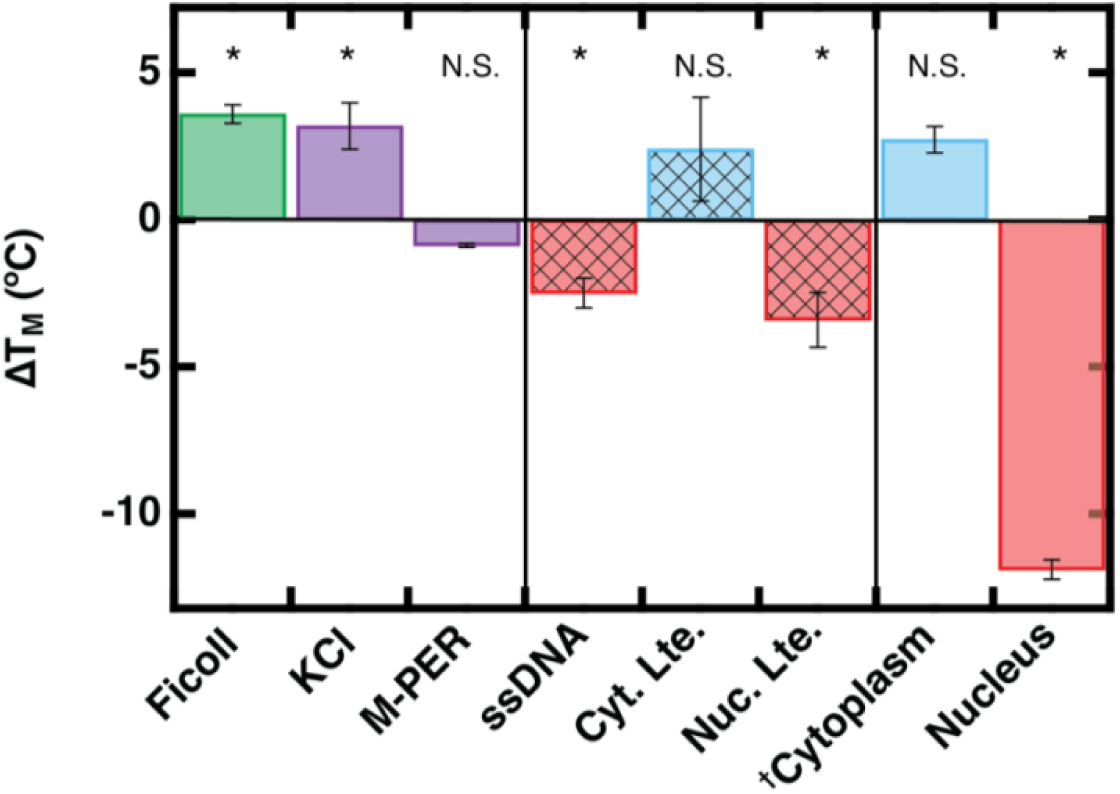
Change in the melting temperature of 0.25 µM FRET-labeled barnase relative to 20 mM sodium phosphate buffer, pH 7.4 in different conditions. The concentrations of in vitro buffer additive are 150 mg/ml Ficoll, 150 mM KCl, 20% M-PER^TM^, 1 mg/ml salmon sperm DNA (ssDNA), ≍1 mg/ml cytoplasmic lysate (Cyt. Lte.), and ≍1 mg/ml nuclear lysate (Nuc. Lte.). Error bars display standard error of at least 3 measurements. *p-value <0.05 and N.S. p-value >0.05. ^†^Reporting T_A_ for the cytoplasm instead of T_M_

Rather, we find that specific protein interactions reproduce cellular trends; while barnase is slightly stabilized in cytoplasmic lysate, it is destabilized in nuclear lysate. This suggests that weak, specific chemical interactions with proteins in either compartment are ultimately responsible for the observed in-cell stability trends. These differences may result from electrostatic or hydrophobic (water-exclusion) attractive interactions between barnase and proteins in each compartment. In general, the larger solvent accessible surface area of an unfolded protein allows for more attractive interactions, thermodynamically favoring the unfolded state over the folded state (Senske et al., 2014). In eukaryotes, cytoplasmic proteins are typically slightly acidic, resulting in an overall negative charge at physiological pH, whereas nuclear proteins span the entire range from acidic to basic, having overall negative, neutral or positive charges (Schwartz et al., 2001). That the stability of barnase is unchanged or slightly increased in the cytosol and cytoplasmic lysate (Figure 5) suggests that attractive and repulsive chemical interactions are offset in these environments. By contrast, the mixture of surface charges in the nucleus results in more attractive interactions with the unfolded state, destabilizing barnase. It is important to note that lysates reproduce the observed cellular trends, but not their magnitudes; this is likely because the protein concentrations in these lysates are significantly lower than in-cell concentrations.

**Figure 5.**
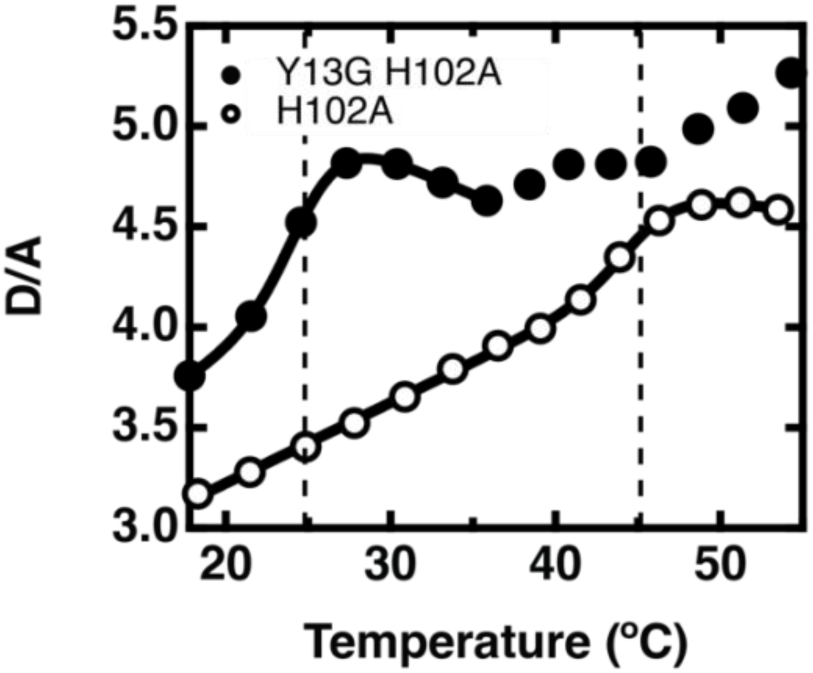
Representative donor/acceptor (D/A) melting curve extracted from FReI of barnase Y13G H102A and barnase H102A overexpressed in BL21 DE3 cells. The data are fit to a two-state denaturation (E1) with dashed vertical lines indicating the melting temperature (T_M_).

### 2.5. Barnase is destabilized and less aggregation prone in bacterial cells

To further investigate the folding of barnase in a more biologically relevant environment we study its folding in BL21 DE3 *E. coli*, a bacterial prokaryote with no compartmentalization: the cytoplasm of *E. coli* is a heterogeneous mixture of highly charged bacterial proteins and nucleic acids (Wennerström et al., 2020; Zimmerman & Trach, 1991). We find that barnase is destabilized in *E. coli* by 15.0 ± 0.7 °C (Figure 5, Table S6). Similar to the nucleus of U2-OS cells, the D/A trends upwards with increasing temperature, unfolding to a stable state that can be observed without aggregation. However, unlike in the nucleus of U2-OS cells, barnase does not typically aggregate within the temperature range of our experiments. These trends hold for pseudo-wild type barnase (barnase H102A), which is destabilized in *E. coli* by 9.5 ± 0.9 °C (Vu et al., 2004) (Figure 5, Table S7). Taken together with the conclusions from U2-OS cells, this suggests that barnase becomes less thermally stable and more conformationally flexible when approaching more native-like environments; however, this increase in conformational flexibility is less likely to be accompanied by unfavorable aggregation that can inhibit function. Proteins have evolved weak interactions to stabilize functional unfolded states in their native environment, like those necessary for the transport of extracellular proteins or to rapidly respond to external stressors. However, such native interactions can be easily disrupted when proteins are moved to a nonnative environment (Mu et al., 2017).

Because it must be inhibited by barstar to protect the host from its ribonuclease activity (Shilova et al., 2021), it is evident that barnase is folded in the *B. amyloliquefaciens* cytosol. However, barnase is an extracellular bacterial protein that is synthesized and secreted by *B. amyloliquefaciens*. In Gram-positive bacteria like *B. amyloliquefaciens,* most secretory proteins, including many degradative enzymes, are unfolded before they are translocated across the cytoplasmic membrane (Sarvas et al., 2004). Indeed, we have previously observed that the outer membrane protein VlsE is more extended and destabilized in the cytosol compared to in vitro (Davis et al., 2020; Guzman et al., 2014). Unfolding could also serve as a functional mechanism to release the tightly bound intracellular inhibitor, barstar, prior to excretion. However, binding is highly irreversible, leading the original discoverers Smeaton and Elliot to hypothesize that (1) the inhibitor is not involved in secretion and (2) active barnase never exists free in the cytoplasm prior to secretion (Smeaton & Elliott, 1967). Instead, they speculated that the inhibitor serves to protect the host organism from enzyme that re-enters the host cytoplasm during stress. In another Gram-positive bacteria, *B. subtilis,* the rate of export of inactive pseudo-wildtype barnase and wildtype barnase in the presence of barstar are comparable (Chen & Nagarajan, 1993). That the rate of transport of wildtype barnase + barstar is only slightly suppressed compared to pseudo-wildtype barnase suggests that the fate of barnase is determined prior to and/or independent of barstar binding (Chen & Nagarajan, 1993). The same study showed that after 90 minutes only 30-40% of barnase is retained inside the cell.

We therefore hypothesize that protein-protein interactions in the host cytosol promote inactive partially folded and extended unfolded conformations of the barnase population that is secreted. This is supported by our in vitro experiments that showed that only cell lysates could fully replicate in cell stability trends (Figure 4). Indeed, Fersht showed that unfolded and partially folded barnase bind molecular chaperones of *E. coli,* GroEL and SecB, with high affinity in vitro (Corrales & Fersht, 1995; Gray & Fersht, 1993; Gray et al., 1993; Zahn et al., 1996). These chaperones decrease the stability of wildtype barnase by 7-10 °C (Zahn et al., 1996), consistent with the destabilization of pseudo-wildtype FRET-labeled barnase measured in *E. coli* (Figure 5). Barnase has also been shown to interact with chaperones overexpressed in eukaryotic cytosols (Wood et al., 2018). Fersht concluded that chaperones have the potential to act in vivo as a folding/transport scaffold, an annealing-machine, and an unfoldase (Zahn et al., 1996). Our data support this mechanism and considered with the limited secretory data available suggest that molecular chaperones must bind barnase early in the folding process to provide another avenue of protection against active cytotoxic barnase. However, this remains to be tested in *B. amyloliquefaciens*.

## 3 Conclusion

We speculate that the decreased thermal stability of barnase in-cell relative to in vitro is representative of increased conformational flexibility in a more functionally relevant context for a ribonuclease, where it must be unfolded prior to excretion. We find that this flexibility is promoted by weak but specific chemical interactions with proteins and nucleic acids in more native-like environments. That barnase does not aggregate upon unfolding in bacterial cells but does aggregate upon unfolding in eukaryotic cells demonstrates that proteins have carefully evolved protein interactions in their native environment. Such interactions may be crucial for functional protein design and interpretation of the evolution of proteomes, which are typically performed on an individual protein or protein-binding partner.

It is not yet possible to computationally predict how protein stability and kinetics change in-cell. Peptides and small proteins offer a path towards bridging this gap between theory and experiment, as experimental data can viably be coupled with all-atom molecular dynamics simulations (Rickard et al., 2020; Rickard et al., 2019). In the future, barnase offers the potential to explore in depth how in vitro protein folding principles translate to the cellular environment. Barnase has an extensive history of being used as a protein folding model, and there exists a large library of mutants designed to probe the contributions of different types of intramolecular interactions to protein folding and stability (Serrano et al., 1992). In vitro barnase has a high affinity for its inhibitor barstar as well as molecular chaperones like GroEL and SecB. Indeed, our work points towards protein-protein interactions as important regulators of barnase folding in cells. Further work remains to untangle how these interactions may be modulated by the crowded cellular environment.

## 4 Materials and Methods

### Protein Expression and Purification

The barnase Y13G H102A gene was inserted into the pET-15b vector for the unlabeled construct and the pDream 2.1 vector for the FRET-labeled construct, both of which contained His tags that could be cleaved by thrombin (GenScript Biotech, Piscataway, NJ). The proteins were expressed and purified from BL21(DE3) competent *E. coli* (New England Biolabs, Ipswich, MA). The cells were grown at 37°C until the OD_600_ reached 0.6, at which point expression was induced with 1 mM isopropyl-β-D-thiogalactopyranoside (Inalco, San Luis Obispo, CA). The induced cells grew overnight at 20°C. The cells were harvested by centrifugation, suspended in lysis buffer, and then lysed by sonication. The resulting lysate was centrifuged and filtered prior to being passed through a Ni-NTA column. Fractions containing protein were confirmed by running SDS-PAGE gels. The resulting fractions were then dialyzed into 20 mM sodium phosphate, pH 7.4. The unlabeled construct further underwent a thrombin cleavage to remove the His tag, using biotinylated thrombin (Millipore Sigma, Burlington, MA). This was also purified using a Ni-NTA column and dialyzed into 20 mM sodium phosphate, pH 7.4. The resulting concentration of the unlabeled construct was determined by absorbance at 280 nm (ε = 26,930 L^-1^ mol^-1^ cm^-1^), and the concentration of the labeled construct was determined by mCherry absorption at 587 nm (ε = 72,000 L^-1^ mol^-1^ cm^-1^). The unlabeled construct was characterized using circular dichroism, to be comparable to the literature (Vu et al., 2004; Vuilleumier et al., 1993), in addition to tryptophan fluorescence (Figure S3). The labeled construct was characterized using FRET.

### Cell Culture and Transfection

Human bone osteosarcoma epithelial cells (U2-OS, ATCC, Manassas, VA) were transfected with the plasmid for either FRET-labeled barnase with a nuclear export sequence or nuclear localization sequence (Table S1). The cells used for transfection were grown in DMEM (Corning, Corning, NY) supplemented with 10% Fetal Bovine Serum (FBS, Gibco, Waltham, MA) and 1% Penicillin-Streptomycin (PS, ThermoFisher Scientific) to a confluency of 70-80%. These cells were split into dishes containing coverslips for subsequent temperature jumps. Initially, these cells are in DMEM with 10% FBS only. To the cells, we added a plasmid-Lipofectamine 2000 (Invitrogen, Waltham, MA) complex prepared in OptiMEM media (Gibco). After 6 hours of incubation at 37°C, the media is exchanged for DMEM with 10% FBS and 1% PS. Temperature jumps were performed after 16 hours of incubation at 37°C. For experiments, the cells were imaged in FluroBrite^TM^ DMEM (Gibco) with 10% FBS.

### Fast Relaxation Imaging

The FReI instrument and methodology have been described elsewhere (Knab & Davis, 2023; Yoo & Davis, 2022). Briefly, the sample is heated by a continuous-wave 2 μm laser (AdValue Photonics, Tucson, AZ). The heating occurs in the form of a step-shaped function, where the temperature is rapidly raised, followed by a 10 second equilibration period at the higher temperature. The sample starts at room temperature and is raised to 50-60°C in ≍2-3°C increments. The temperature at each step is determined based on calibration with mCherry, which has a well-established temperature-dependent quantum yield. For in-cell FRET measurements, AcGFP1 was excited with blue light selected from a white light LED source (X-Cite Mini +, Excelitas Technology Corporation, Waltham, MA), using a ET480/40x bandpass filter (Chroma, Bellows Falls, VT) and a FF509-FDi01 dichroic mirror (Semrock, West Henrietta, NY). To avoid detection of excitation light, an ET510lp long pass filter (Chroma) selects for emission light. The emission is then separated into green and red light by T510lpxrxt-UF3 and ZT594rdc-UF3 dichroic mirrors (Chroma). The collected light is detected by a CMOS camera (Moment, Teledyne, Thousand Oaks, CA) with an integration time of 200 ms for cells.

### Buffer Preparation

All in vitro measurements were performed in 20 mM sodium phosphate, pH 7.4. All in vitro tryptophan melts for the unlabeled protein were performed at a concentration of 10 μM, and all in vitro FRET melts for the labeled protein were performed at a concentration of 0.25 μM. Table S5 contains data for all in vitro melts.

Ficoll PM 70 (Sigma-Aldrich, St. Louis, MO) and M-PER^TM^ (ThermoFisher Scientific, Waltham, MA) melts were performed using tryptophan fluorescence of the unlabeled protein. The Ficoll solutions were prepared by first creating a 400 mg/mL stock solution in 20 mM sodium phosphate, pH 7.4 and then diluting to the final working concentration. The stock solution was left in a 55°C water bath overnight. Working solutions were further sonicated for 15 minutes prior to addition of the protein.

Melts in salmon sperm DNA (Invitrogen, Waltham, MA) were performed using the FRET-labeled protein in 20 mM sodium phosphate, pH 7.4. The final concentration of salmon sperm DNA was 1 mg/mL.

Melts in cytoplasmic and nuclear lysate were performed using the FRET-labeled protein. The lysates were prepared from U2-OS cells at 70-80% confluency, following the manufacturer’s protocol for the NE-PER^TM^ Nuclear and Cytoplasmic Extraction Reagents (Thermo Scientific, Waltham, MA). The lysates were dialyzed into 20 mM sodium phosphate, pH 7.4 for experiments. Control experiments in the extraction reagents alone confirm that any trace extraction reagent cannot account for observations in lysates (Figure S4). The concentration of protein in the extracts was determined using a Bradford assay. Concentrations were in the range of ≍600-4000 μg/mL, with generally lower yields for nuclear proteins.

### Fluorescence Melts

Fluorescence melts were performed on a Jasco FP-8500 (Easton, MD). For tryptophan melts, the excitation wavelength was 280 nm and emission was monitored across 275 to 450 nm. For FRET melts, the excitation wavelength was 475 nm and emission was monitored across 470 to 700 nm. Melts were performed from 20 to 71 °C in 3 °C increments, with an equilibration period of 3 minutes at each temperature.

### Preparation of *E. Coli*

The plasmid for FRET-labeled barnase was transformed in BL21(DE3) *E. coli*. The cells were grown at 37°C until the OD_600_ reached 0.6, at which point expression was induced with 1 mM isopropyl-β-D-thiogalactopyranoside (Inalco, San Luis Obispo, CA). The induced cells grew overnight at 20°C. *E. coli* were prepared for temperature jumps by initially spinning down 10-mL of culture at 4000 rpm for 10 minutes at 4 °C. The resulting pellet was then resuspended in 10-mL of PBS (Corning, Corning, NY) that had been refrigerated overnight. The PBS-suspended cells were then centrifuged at 4000 rpm for 10 minutes at 4 °C, and this process was repeated for a total of 3 times. After the last wash, the pellet was resuspended in 200 μL of PBS. This was used for temperature jumps, with an exposure time of 100-200 ms.

### Analysis of Thermodynamic Data

For tryptophan melts, the intensity at the wavelength of maximum emission was plotted against temperature. Given that the large linear baselines resulting from the temperature-dependent quantum yields of the fluorophores can obscure the melting transition for small proteins, we removed the sloping baselines by subtracting the acceptor intensity scaled by the D/A at the lowest temperature from the donor intensity for in vitro FRET melts (Girdhar et al., 2011; Knab & Davis, 2023). An artifact of determining thermodynamics this was is that the absolute value of the melting temperature is inaccurate, however, all data corrected at the same temperature can be compared so that the change in melting temperatures are accurate (ΔT_m_). Thus, for in vitro FRET melts, the donor – scaled acceptor intensity (D-aA) was plotted against temperature (Table S4-S5). For in-cell FRET melts and temperature jumps, the donor/acceptor ratio (D/A) was plotted against temperature (Table S2-S3, S6-S7). The resulting melting curve was fit to the following two-state denaturation equation:

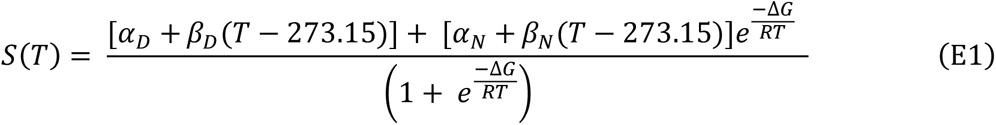

Where N and D denote the native and denatured states, *α_N_* and *α_D_* are the signals from each state at 0°C, *β_N_* and *β_D_* are the slopes of the baselines for each state, and ΔG is the free energy of folding.

## 5 Supplementary Material Description

Supplementary materials include a representative circular dichroism and fluorescence melt of unlabeled barnase, representative temperature-dependent behavior of aggregated in-cell populations, melts in cytoplasmic and nuclear extraction reagents, a table of protein sequences, and tables of melting temperatures collected in vitro and in living cells.

## 6 Acknowledgements

This work was supported by National Institutes of Health (NIH) grant R35 GM151146. U.T. was partially supported by the NIH under Biophysics Training Grant T32 GM008283.

## 7. Conflict of Interests

The authors declare no conflict of interest

## Notes

### Competing Interest Statement

The authors have declared no competing interest.

### Summary of Updates

Figure 4 revised; supplemental file updated

## References

Alemany, A., Rey-Serra, B., Frutos, S., Cecconi, C., & Ritort, F. (2016). Mechanical Folding and Unfolding of Protein Barnase at the Single-Molecule Level. Biophysical Journal, 110(1), 63–74. doi: 10.1016/j.bpj.2015.11.015

Ames, A., & Nesbett, F. B. (1966). Intracellular and extracellular compartments of mammalian central nervous tissue. The Journal of Physiology, 184(1), 215–238.

Benton, L. A., Smith, A. E., Young, G. B., & Pielak, G. J. (2012). Unexpected Effects of Macromolecular Crowding on Protein Stability. Biochemistry, 51(49), 9773–9775. doi: 10.1021/bi300909q

Brocchieri, L., & Karlin, S. (2005). Protein length in eukaryotic and prokaryotic proteomes. Nucleic Acids Research, 33(10), 3390–3400. doi: 10.1093/nar/gki615

Bye, J. W., Baxter, N. J., Hounslow, A. M., Falconer, R. J., & Williamson, M. P. (2016). Molecular Mechanism for the Hofmeister Effect Derived from NMR and DSC Measurements on Barnase. ACS Omega, 1(4), 669–679. doi: 10.1021/acsomega.6b00223

Chen, M., & Nagarajan, V. (1993). The roles of signal peptide and mature protein in RNase (barnase) export from Bacillus subtilis. Molecular & General Genetics: MGG, 239(3), 409–415. doi: 10.1007/BF00276939

Chen, W. W., Freinkman, E., Wang, T., Birsoy, K., & Sabatini, D. M. (2016). Absolute Quantification of Matrix Metabolites Reveals the Dynamics of Mitochondrial Metabolism. Cell, 166(5), 1324–1337.e11. doi: 10.1016/j.cell.2016.07.040

Christiansen, A., Wang, Q., Samiotakis, A., Cheung, M. S., & Wittung-Stafshede, P. (2010). Factors Defining Effects of Macromolecular Crowding on Protein Stability: An in Vitro/in Silico Case Study Using Cytochrome c. Biochemistry, 49(31), 6519–6530. doi: 10.1021/bi100578x

Chu, I.-T., Hutcheson, B. O., Malsch, H. R., & Pielak, G. J. (2023). Macromolecular Crowding by Polyethylene Glycol Reduces Protein Breathing. The Journal of Physical Chemistry Letters, 14(10), 2599–2605. doi: 10.1021/acs.jpclett.3c00271

Corrales, F. J., & Fersht, A. R. (1995). The folding of GroEL-bound barnase as a model for chaperonin-mediated protein folding. Proceedings of the National Academy of Sciences of the United States of America, 92(12), 5326–5330. doi: 10.1073/pnas.92.12.5326

Danielsson, J., Mu, X., Lang, L., Wang, H., Binolfi, A., Theillet, F.-X., … Oliveberg, M. (2015). Thermodynamics of protein destabilization in live cells. Proceedings of the National Academy of Sciences, 112(40), 12402–12407. doi: 10.1073/pnas.1511308112

Davis, C. M., Deutsch, J., & Gruebele, M. (2020). An in vitro mimic of in-cell solvation for protein folding studies. Protein Science: A Publication of the Protein Society, 29(4), 1060–1068. doi: 10.1002/pro.3833

Davis, C. M., & Gruebele, M. (2018). Non-Steric Interactions Predict the Trend and Steric Interactions the Offset of Protein Stability in Cells. ChemPhysChem, 19(18), 2290–2294. doi: 10.1002/cphc.201800534

Davis, C. M., Gruebele, M., & Sukenik, S. (2018). How does solvation in the cell affect protein folding and binding? Current Opinion in Structural Biology, 48, 23–29. doi: 10.1016/j.sbi.2017.09.003

Dhar, A., Girdhar, K., Singh, D., Gelman, H., Ebbinghaus, S., & Gruebele, M. (2011). Protein Stability and Folding Kinetics in the Nucleus and Endoplasmic Reticulum of Eucaryotic Cells. Biophysical Journal, 101(2), 421–430. doi: 10.1016/j.bpj.2011.05.071

Ebbinghaus, S., Dhar, A., McDonald, J. D., & Gruebele, M. (2010). Protein folding stability and dynamics imaged in a living cell. Nature Methods, 7(4), 319–323. doi: 10.1038/nmeth.1435

Feng, R., Gruebele, M., & Davis, C. M. (2019). Quantifying protein dynamics and stability in a living organism. Nature Communications, 10(1), 1179. doi: 10.1038/s41467-019-09088-y

Fersht, A. R. (1993). Protein folding and stability: The pathway of folding of barnase. FEBS Letters, 325(1–2), 5–16. doi: 10.1016/0014-5793(93)81405-O

Fersht, A. R. (2000). A kinetically significant intermediate in the folding of barnase. Proceedings of the National Academy of Sciences, 97(26), 14121–14126. doi: 10.1073/pnas.260502597

Girdhar, K., Scott, G., Chemla, Y. R., & Gruebele, M. (2011). Better biomolecule thermodynamics from kinetics. The Journal of Chemical Physics, 135(1), 015102. doi: 10.1063/1.3607605

Gray, Tamara E., & Fersht, A. R. (1993). Refolding of Barnase in the Presence of GroE. Journal of Molecular Biology, 232(4), 1197–1207. doi: 10.1006/jmbi.1993.1471

Gray, T.E., Eder, J., Bycroft, M., Day, A. G., & Fersht, A. R. (1993). Refolding of barnase mutants and pro-barnase in the presence and absence of GroEL. The EMBO Journal, 12(11), 4145–4150. doi: 10.1002/j.1460-2075.1993.tb06098.x

Guigas, G., Kalla, C., & Weiss, M. (2007). The degree of macromolecular crowding in the cytoplasm and nucleoplasm of mammalian cells is conserved. FEBS Letters, 581(26), 5094–5098. doi: 10.1016/j.febslet.2007.09.054

Guzman, I., Gelman, H., Tai, J., & Gruebele, M. (2014). The Extracellular Protein VlsE Is Destabilized Inside Cells. Journal of Molecular Biology, 426(1), 11–20. doi: 10.1016/j.jmb.2013.08.024

Hartley, R. W. (1989). Barnase and barstar: Two small proteins to fold and fit together. Trends in Biochemical Sciences, 14(11), 450–454. doi: 10.1016/0968-0004(89)90104-7

Hartley, R. W. (1993). Directed mutagenesis and barnase-barstar recognition. Biochemistry, 32(23), 5978–5984. doi: 10.1021/bi00074a008

Johnson, C. M., & Fersht, A. R. (1995). Protein Stability as a Function of Denaturant Concentration: The Thermal Stability of Barnase in the Presence of Urea. Biochemistry, 34(20), 6795–6804. doi: 10.1021/bi00020a026

Kellermayer, M. (1981). Soluble and “Loosely Bound” Nuclear Proteins in Regulation of the Ionic Environment in Living Cell Nuclei. In H. G. Schweiger (Ed.), International Cell Biology 1980–1981 (pp. 915–924). Berlin, Heidelberg: Springer Berlin Heidelberg. doi: 10.1007/978-3-642-67916-2_103

Khan, F., Chuang, J. I., Gianni, S., & Fersht, A. R. (2003). The Kinetic Pathway of Folding of Barnase. Journal of Molecular Biology, 333(1), 169–186. doi: 10.1016/j.jmb.2003.08.024

Kippen, A. D., Sancho, J., & Fersht, A. R. (1994). Folding of Barnase in Parts. Biochemistry, 33(12), 3778–3786. doi: 10.1021/bi00178a039

Kırlı, K., Karaca, S., Dehne, H. J., Samwer, M., Pan, K. T., Lenz, C., … Görlich, D. (2015). A deep proteomics perspective on CRM1-mediated nuclear export and nucleocytoplasmic partitioning. eLife, 4, e11466. doi: 10.7554/eLife.11466

Knab, E., & Davis, C. M. (2023). Chemical interactions modulate λ6-85 stability in cells. Protein Science, 32(7), e4698. doi: 10.1002/pro.4698

Lee, L.-P., & Tidor, B. (2001). Barstar is electrostatically optimized for tight binding to barnase. Nature Structural Biology, 8(1), 73–76. doi: 10.1038/83082

McConkey, E. H. (1982). Molecular evolution, intracellular organization, and the quinary structure of proteins. Proceedings of the National Academy of Sciences, 79(10), 3236–3240. doi: 10.1073/pnas.79.10.3236

Meiering, E. M., Serrano, L., & Fersht, A. R. (1992). Effect of active site residues in barnase on activity and stability. Journal of Molecular Biology, 225(3), 585–589. doi: 10.1016/0022-2836(92)90387-Y

Monteith, W. B., Cohen, R. D., Smith, A. E., Guzman-Cisneros, E., & Pielak, G. J. (2015). Quinary structure modulates protein stability in cells. Proceedings of the National Academy of Sciences, 112(6), 1739–1742. doi: 10.1073/pnas.1417415112

Mu, X., Choi, S., Lang, L., Mowray, D., Dokholyan, N. V., Danielsson, J., & Oliveberg, M. (2017). Physicochemical code for quinary protein interactions in Escherichia coli. Proceedings of the National Academy of Sciences, 114(23), E4556–E4563. doi: 10.1073/pnas.1621227114

Paine, P. L., Pearson, T. W., Tluczek, L. J. M., & Horowitz, S. B. (1981). Nuclear sodium and potassium. Nature, 291(5812), 258–261. doi: 10.1038/291258a0

Qi, H. W., Nakka, P., Chen, C., & Radhakrishnan, M. L. (2014). The Effect of Macromolecular Crowding on the Electrostatic Component of Barnase–Barstar Binding: A Computational, Implicit Solvent-Based Study. PLOS ONE, 9(6), e98618. doi: 10.1371/journal.pone.0098618

Raeburn, C. B., Ormsby, A. R., Cox, D., Gerak, C. A., Makhoul, C., Moily, N. S., … Hatters, D. M. (2022). A biosensor of protein foldedness identifies increased “holdase” activity of chaperones in the nucleus following increased cytosolic protein aggregation. Journal of Biological Chemistry, 298(8). doi: 10.1016/j.jbc.2022.102158

Rickard, M. M., Zhang, Y., Pogorelov, T. V., & Gruebele, M. (2020). Crowding, Sticking, and Partial Folding of GTT WW Domain in a Small Cytoplasm Model. The Journal of Physical Chemistry B, 124(23), 4732–4740. doi: 10.1021/acs.jpcb.0c02536

Rickard, Meredith M., Zhang, Y., Gruebele, M., & Pogorelov, T. V. (2019). In-Cell Protein–Protein Contacts: Transient Interactions in the Crowd. The Journal of Physical Chemistry Letters, 10(18), 5667–5673. doi: 10.1021/acs.jpclett.9b01556

Ridgway, D., Broderick, G., Lopez-Campistrous, A., Ru’aini, M., Winter, P., Hamilton, M., … Ellison, M. J. (2008). Coarse-Grained Molecular Simulation of Diffusion and Reaction Kinetics in a Crowded Virtual Cytoplasm. Biophysical Journal, 94(10), 3748–3759. doi: 10.1529/biophysj.107.116053

Sarvas, M., Harwood, C. R., Bron, S., & van Dijl, J. M. (2004). Post-translocational folding of secretory proteins in Gram-positive bacteria. Biochimica et Biophysica Acta (BBA) - Molecular Cell Research, 1694(1), 311–327. doi: 10.1016/j.bbamcr.2004.04.009

Schreiber, G., Buckle, A. M., & Fersht, A. R. (1994). Stability and function: Two constraints in the evolution of barstar and other proteins. Structure, 2(10), 945– 951. doi: 10.1016/S0969-2126(94)00096-4

Schwartz, R., Ting, C. S., & King, J. (2001). Whole Proteome pI Values Correlate with Subcellular Localizations of Proteins for Organisms within the Three Domains of Life. Genome Research, 11(5), 703–709. doi: 10.1101/gr.158701

Senske, M., Törk, L., Born, B., Havenith, M., Herrmann, C., & Ebbinghaus, S. (2014). Protein Stabilization by Macromolecular Crowding through Enthalpy Rather Than Entropy. Journal of the American Chemical Society, 136(25), 9036–9041. doi: 10.1021/ja503205y

Serrano, L., Kellis, J. T., Cann, P., Matouschek, A., & Fersht, A. R. (1992). The folding of an enzyme: II. Substructure of barnase and the contribution of different interactions to protein stability. Journal of Molecular Biology, 224(3), 783–804. doi: 10.1016/0022-2836(92)90562-X

Shilova, O., Kotelnikova, P., Proshkina, G., Shramova, E., & Deyev, S. (2021). Barnase-Barstar Pair: Contemporary Application in Cancer Research and Nanotechnology. Molecules, 26(22), 6785. doi: 10.3390/molecules26226785

Sieg, J. P., McKinley, L. N., Huot, M. J., Yennawar, N. H., & Bevilacqua, P. C. (2022). The Metabolome Weakens RNA Thermodynamic Stability and Strengthens RNA Chemical Stability. Biochemistry, 61(22), 2579–2591. doi: 10.1021/acs.biochem.2c00488

Smeaton, J. R., & Elliott, W. H. (1967). Isolation and properties of a specific bacterial ribonuclease inhibitor. Biochimica et Biophysica Acta (BBA) - Nucleic Acids and Protein Synthesis, 145(3), 547–560. doi: 10.1016/0005-2787(67)90115-3

Smith, A. E., Zhou, L. Z., Gorensek, A. H., Senske, M., & Pielak, G. J. (2016). In-cell thermodynamics and a new role for protein surfaces. Proceedings of the National Academy of Sciences, 113(7), 1725–1730. doi: 10.1073/pnas.1518620113

Spaar, A., Dammer, C., Gabdoulline, R. R., Wade, R. C., & Helms, V. (2006). Diffusional Encounter of Barnase and Barstar. Biophysical Journal, 90(6), 1913– 1924. doi: 10.1529/biophysj.105.075507

Steinberg, R., & Koch, H.-G. (2021). The largely unexplored biology of small proteins in pro- and eukaryotes. The FEBS Journal, 288(24), 7002–7024. doi: 10.1111/febs.15845

Storz, G., Wolf, Y. I., & Ramamurthi, K. S. (2014). Small Proteins Can No Longer Be Ignored. Annual Review of Biochemistry, 83(Volume 83, 2014), 753–777. doi: 10.1146/annurev-biochem-070611-102400

Tai, J., Dave, K., Hahn, V., Guzman, I., & Gruebele, M. (2016). Subcellular modulation of protein VlsE stability and folding kinetics. FEBS Letters, 590(10), 1409–1416. doi: 10.1002/1873-3468.12193

The PyMOL Molecular Graphics System. (n.d.). Schrödinger, LLC.

Thole, J. F., Fadero, T. C., Bonin, J. P., Stadmiller, S. S., Giudice, J. A., & Pielak, G. J. (2021). Danio rerio Oocytes for Eukaryotic In-Cell NMR. Biochemistry, 60(6), 451–459. doi: 10.1021/acs.biochem.0c00922

Trevitt, C. R., Yashwanth Kumar, D. R., Fowler, N. J., & Williamson, M. P. (2024). Interactions between the protein barnase and co-solutes studied by NMR. Communications Chemistry, 7(1), 1–15. doi: 10.1038/s42004-024-01127-0

Ulyanova, V., Vershinina, V., & Ilinskaya, O. (2011). Barnase and binase: Twins with distinct fates. The FEBS Journal, 278(19), 3633–3643. doi: 10.1111/j.1742-4658.2011.08294.x

van der Zanden, S. Y., Jongsma, M. L. M., Neefjes, A. C. M., Berlin, I., & Neefjes, J. (2023). Maintaining soluble protein homeostasis between nuclear and cytoplasmic compartments across mitosis. Trends in Cell Biology, 33(1), 18–29. doi: 10.1016/j.tcb.2022.06.002

Vu, N.-D., Feng, H., & Bai, Y. (2004). The Folding Pathway of Barnase: The Rate-Limiting Transition State and a Hidden Intermediate under Native Conditions. Biochemistry, 43(12), 3346–3356. doi: 10.1021/bi0362267

Vuilleumier, S., Sancho, J., Loewenthal, R., & Fersht, A. R. (1993). Circular dichroism studies of barnase and its mutants: Characterization of the contribution of aromatic side chains. Biochemistry, 32(39), 10303–10313. doi: 10.1021/bi00090a005

Wennerström, H., Vallina Estrada, E., Danielsson, J., & Oliveberg, M. (2020). Colloidal stability of the living cell. Proceedings of the National Academy of Sciences, 117(19), 10113–10121. doi: 10.1073/pnas.1914599117

Wood, R. J., Ormsby, A. R., Radwan, M., Cox, D., Sharma, A., Vöpel, T., … Hatters, D. M. (2018). A biosensor-based framework to measure latent proteostasis capacity. Nature Communications, 9(1), 287. doi: 10.1038/s41467-017-02562-5

Yoo, H., & Davis, C. M. (2022). An in Vitro Cytomimetic of In-Cell RNA Folding. ChemBioChem, 23(20), e202200406. doi: 10.1002/cbic.202200406

Zahn, R., Perrett, S., & Fersht, A. R. (1996). Conformational states bound by the molecular chaperones GroEL and secB: A hidden unfolding (annealing) activity. Journal of Molecular Biology, 261(1), 43–61. doi: 10.1006/jmbi.1996.0440

Zhou, H.-X., Rivas, G., & Minton, A. P. (2008). Macromolecular Crowding and Confinement: Biochemical, Biophysical, and Potential Physiological Consequences*. Annual Review of Biophysics, 37(Volume 37, 2008), 375–397. doi: 10.1146/annurev.biophys.37.032807.125817

Zimmerman, S. B., & Trach, S. O. (1991). Estimation of macromolecule concentrations and excluded volume effects for the cytoplasm of Escherichia coli. Journal of Molecular Biology, 222(3), 599–620. doi: 10.1016/0022-2836(91)90499-v

