## Supplementary Information for "Weak, specific chemical interactions dictate barnase stability in diverse cellular environments"

### Results and Discussion:

**Figure S1**

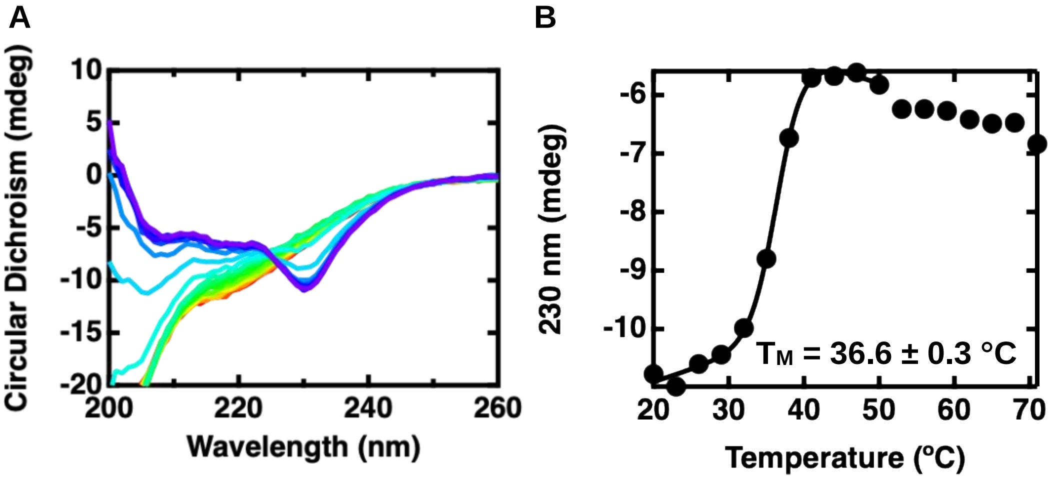


**Figure S1.** (A) Representative circular dichroism (CD) spectra for 10 μM barnase in 20 mM sodium phosphate, pH 7.4 between 20 and 71 °C in 3 °C increments. (B) CD signal monitored at 230 nm, a characteristic peak for barnase (Vuilleumier et al., 1993), as a function of temperature and fit to E1.

**Figure S2**

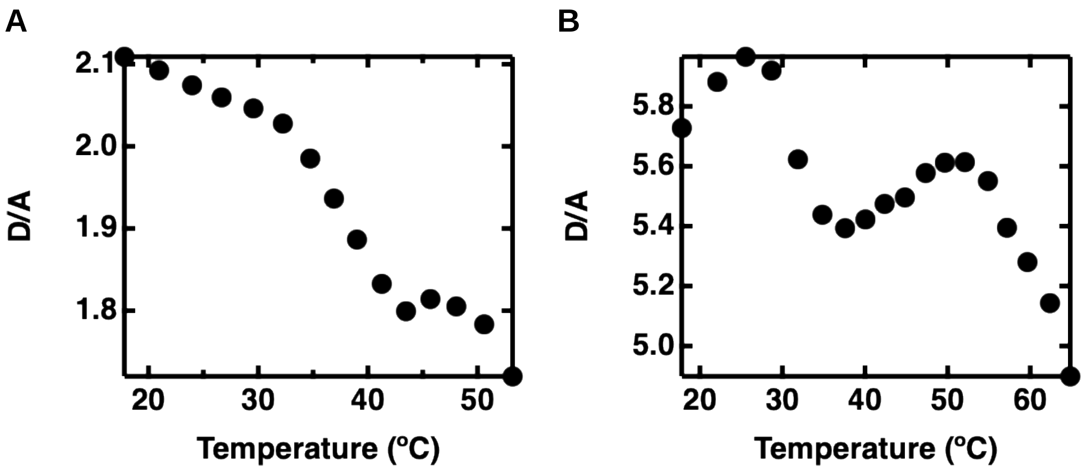


**Figure S2.** Example temperature jump data for unfolded and aggregated populations in the nucleus for (A) a pre-aggregated population with a lower than average initial D/A and (B) an unfolded population with a higher than average initial D/A.

**Figure S3**

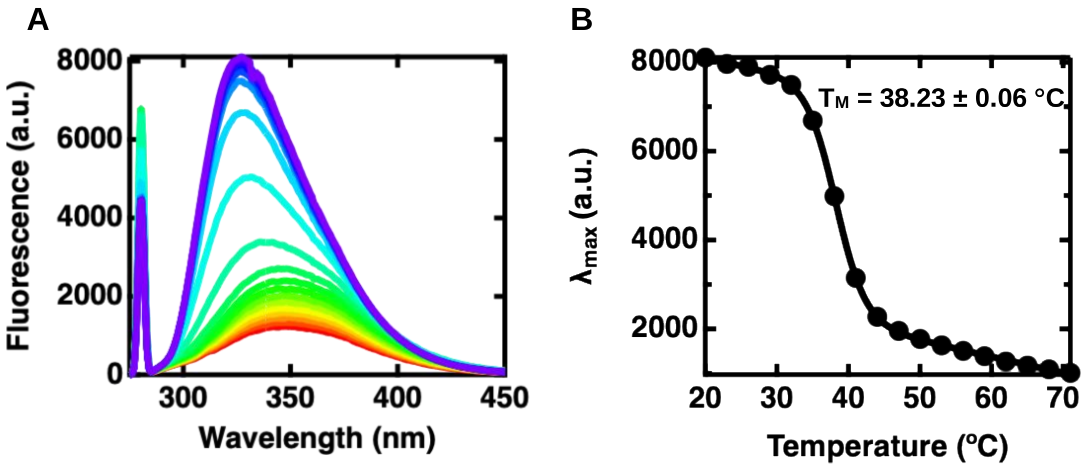


**Figure S3.** (A) Representative tryptophan fluorescence spectra for 10 μM barnase in 20 mM sodium phosphate, pH 7.4 between 20 and 71 °C in 3 °C increments. (B) Tryptophan fluorescence intensity monitored at the wavelength of maximum emission as a function of temperature and fit to E1.

### Figure S4

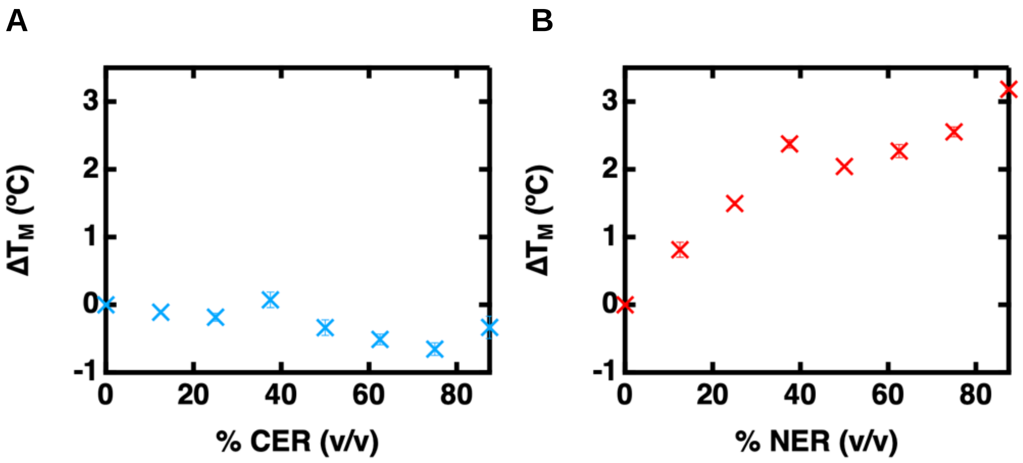


**Figure S4.** Change in the melting temperature of 10 μM unlabeled barnase in increasing concentrations of (A) cytoplasmic extraction reagent and (B) nuclear extraction reagent

**Table S1.** Protein sequences used in the paper, where underlined text is the sequence for barnase, and bold text is either a nuclear export or nuclear localization sequence (Raeburn et al., 2022)

| Protein | Sequence |
|---|---|
| Unlabeled Barnase | MGSSHHHHHH SSGLVPRGSH MAQVINTFDG VADGLQTYHK<br>LPDNYITKSE AQALGWVASK GNLADVAPGK SIGGDIFSNR<br>EGKLPGKSGR TWREADINYT SGFRNSDRIL YSSDWLIYKT<br>TDAYQTFTKI R |
| FRET-Labeled Barnase | MDYKDDDDKG SGSSHHHHHH SSGLVPRGSM VSKGAELFTG<br>IVPILIELNG DVNGHKFSVS GEGEGDATYG KLTLKFICTT<br>GKLPVPWPTL VTTLSYGVQC FSRYPDHMKQ HDFFKSAMPE<br>GYIQERTIFF EDDGNYKSRA EVKFEGDTLV NRIELTGTDF<br>KEDGNILGNK MEYNYNAHNV YIMTDKAKNG IKVNFKIRHN<br>IEDGSVQLAD HYQQNTPIGD GPVLLPDNHY LSTQSALSKD<br>PNEKRDHMIY FGFVTAAAIT HGMDELYKHM AQVINTFDGV<br>ADGLQTYHKL PDNYITKSEA QALGWVASKG NLADVAPGKS<br>IGGDIFSNRE GKLPGKSGRT WREADINYTS GFRNSDRILY<br>SSDWLIYKTT DAYQTFTKIR KLMVSKGEED NMAIIKEFMR<br>FKVHMEGSVN GHEFEIEGEG EGRPYEGTQT AKLKVTKGGP<br>LPFAWDILSP QFMYGSKAYV KHPADIPDYL KLSFPEGFKW<br>ERVMNFEDGG VVTVTQDSSL QDGEFIYKVK LRGTNFPSDG<br>PVMQKKTMGW EASSERMYPE DGALKGEIKQ RLKLKDGGHY<br>DAEVKTTYKA KKPVQLPGAY NVNIKLDITS HNEDYTIVEQ<br>YERAEGRHST GGMDELYK |

| | |
|---|---|
| FRET-Labeled Barnase with **Nuclear Export Sequence** | MDYKDDDDKG SGSSHHHHHH SSGLVPRGSM VSKGAELFTG<br>IVPILIELNG DVNGHKFSVS GEGEGDATYG KLTLKFICTT<br>GKLPVPWPTL VTTLSYGVQC FSRYPDHMKQ HDFFKSAMPE<br>GYIQERTIFF EDDGNYKSRA EVKFEGDTLV NRIELTGTDF<br>KEDGNILGNK MEYNYNAHNV YIMTDKAKNG IKVNFKIRHN<br>IEDGSVQLAD HYQQNTPIGD GPVLLPDNHY LSTQSALSKD<br>PNEKRDHMIY FGFVTAAAIT HGMDELYKHM AQVINTFDGV<br>ADGLQTYHKL PDNYITKSEA QALGWVASKG NLADVAPGKS<br>IGGDIFSNRE GKLPGKSGRT WREADINYTS GFRNSDRILY<br>SSDWLIYKTT DAYQTFTKIR **LQKKLEELEL DE**KLMVSKGE<br>EDNMAIIKEF MRFKVHMEGS VNGHEFEIEG EGEGRPYEGT<br>QTAKLKVTKG GPLPFAWDIL SPQFMYGSKA YVKHPADIPD<br>YLKLSFPEGF KWERVMNFED GGVVTVTQDS SLQDGEFIYK<br>VKLRGTNFPS DGPVMQKKTM GWEASSERMY PEDGALKGEI<br>KQRLKLKDGG HYDAEVKTTY KAKKPVQLPG AYNVNIKLDI<br>TSHNEDYTIV EQYERAEGRH STGGMDELYK |
| FRET-Labeled Barnase with **Nuclear Localization Sequence** | MDYKDDDDKG SGSSHHHHHH SSGLVPRGSM VSKGAELFTG<br>IVPILIELNG DVNGHKFSVS GEGEGDATYG KLTLKFICTT<br>GKLPVPWPTL VTTLSYGVQC FSRYPDHMKQ HDFFKSAMPE<br>GYIQERTIFF EDDGNYKSRA EVKFEGDTLV NRIELTGTDF<br>KEDGNILGNK MEYNYNAHNV YIMTDKAKNG IKVNFKIRHN<br>IEDGSVQLAD HYQQNTPIGD GPVLLPDNHY LSTQSALSKD<br>PNEKRDHMIY FGFVTAAAIT HGMDELYKHM **MCGGGPKKKR**<br>**KVED**AQVINT FDGVADGLQT YHKLPDNYIT KSEAQALGWV<br>ASKGNLADVA PGKSIGGDIF SNREGKLPGK SGRTWREADI<br>NYTSGFRNSD RILYSSDWLI YKTTDAYQTF TKIRKLMVSK<br>GEEDNMAIIK EFMRFKVHME GSVNGHEFEI EGEGEGRPYE<br>GTQTAKLKVT KGGPLPFAWD ILSPQFMYGS KAYVKHPADI<br>PDYLKLSFPE GFKWERVMNF EDGGVVTVTQ DSSLQDGEFI<br>YKVKLRGTNF PSDGPVMQKK TMGWEASSER MYPEDGALKG<br>EIKQRLKLKD GGHYDAEVKT TYKAKKPVQL PGAYNVNIKL<br>DITSHNEDYT IVEQYERAEG RHSTGGMDEL YK |

**Table S2.** $T_A$ values for cytoplasm of U2-OS cells extracted from fitting D/A vs. temperature

| U2-OS Cell Number | $T_A$ (°C) |
|---|---|
| 1 | 40.4 ± 0.4 |
| 2 | 37.4 ± 0.4 |
| 3 | 38.0 ± 0.5 |
| 4 | 41.5 ± 0.4 |
| 5 | 39.2 ± 0.5 |
| 6 | 38.9 ± 0.5 |
| 7 | 40.3 ± 0.4 |

| | |
|---|---|
| 8 | 42 ± 2 |
| 9 | 37.1 ± 0.6 |
| 10 | 36.3 ± 0.2 |
| 11 | 39.7 ± 0.8 |
| 12 | 45.4 ± 0.9 |
| 13 | 44.3 ± 0.8 |
| 14 | 42.0 ± 0.4 |
| 15 | 42.3 ± 0.7 |
| 16 | 41.7 ± 0.5 |
| 17 | 39.4 ± 0.3 |
| 18 | 41 ± 1 |
| 19 | 43 ± 1 |
| 20 | 41.9 ± 0.5 |
| 21 | 44 ± 2 |
| 22 | 43.0 ± 0.6 |
| 23 | 45.1 ± 0.6 |
| 24 | 42.3 ± 0.2 |
| 25 | 41.4 ± 0.2 |
| 26 | 46.3 ± 0.4 |
| 27 | 29.9 ± 0.2 |
| 28 | 42.3 ± 0.9 |
| 29 | 40.2 ± 0.4 |
| 30 | 41.7 ± 0.6 |
| 31 | 38.4 ± 0.3 |
| 32 | 41.6 ± 0.4 |
| 33 | 38.4 ± 0.3 |
| 34 | 41.1 ± 0.4 |
| 35 | 44.2 ± 0.6 |
| 36 | 42.7 ± 0.7 |
| 37 | 35.8 ± 0.2 |
| 38 | 37.1 ± 0.3 |
| 39 | 40 ± 2 |
| 40 | 45.7 ± 0.7 |
| 41 | 48.7 ± 0.3 |
| 42 | 40.9 ± 0.6 |
| 43 | 43.4 ± 0.6 |
| 44 | 46.3 ± 0.9 |
| 45 | 38.9 ± 0.7 |
| 46 | 30.9 ± 0.2 |

| | |
|---|---|
| 47 | 37 ± 2 |
| 48 | 44.7 ± 0.6 |
| 49 | 47 ± 2 |
| 50 | 42.6 ± 0.7 |
| 51 | 47.7 ± 0.4 |
| 52 | 38.3 ± 0.4 |
| 53 | 42.2 ± 0.7 |
| 54 | 39 ± 1 |
| 55 | 50.4 ± 0.4 |
| 56 | 38.2 ± 0.8 |
| 57 | 42.5 ± 0.5 |
| 58 | 45 ± 3 |
| 59 | 39.3 ± 0.2 |
| 60 | 48.0 ± 0.3 |
| 61 | 45.7 ± 0.3 |
| 62 | 43.3 ± 0.5 |
| 63 | 41 ± 2 |
| 64 | 40.7 ± 0.5 |
| 65 | 42.9 ± 0.5 |
| 66 | 44.6 ± 0.5 |
| 67 | 32.6 ± 0.5 |
| 68 | 42.5 ± 0.5 |
| 69 | 42 ± 1 |
| 70 | 40.8 ± 0.8 |
| **AVERAGE** | **41.4 ± 0.5** |
| **P-Value** | 0.155 |

**Table S3.** $T_M$ values for nucleus of U2-OS cells extracted from fitting D/A vs. temperature

| **U2-OS Cell Number** | **$T_M$ (ºC)** |
|---|---|
| 1 | 26 ± 1 |
| 2 | 26.0 ± 0.6 |
| 3 | 26.6 ± 0.6 |
| 4 | 24 ± 2 |
| 5 | 24 ± 1 |
| 6 | 30 ± 3 |
| 7 | 27 ± 1 |

| | |
|---|---|
| 8 | 28.4 ± 0.3 |
| 9 | 23.0 ± 0.8 |
| 10 | 27 ± 1 |
| 11 | 24 ± 1 |
| 12 | 23.1 ± 0.8 |
| 13 | 24.2 ± 0.6 |
| 14 | 26.1 ± 0.8 |
| 15 | 25.3 ± 0.6 |
| 16 | 26 ± 1 |
| 17 | 26.2 ± 0.4 |
| 18 | 30.0 ± 0.4 |
| 19 | 26.8 ± 0.8 |
| 20 | 26.6 ± 0.7 |
| 21 | 26.4 ± 0.5 |
| 22 | 23.4 ± 0.8 |
| 23 | 26.8 ± 0.4 |
| 24 | 27.9 ± 0.6 |
| 25 | 29.1 ± 0.5 |
| 26 | 28.7 ± 0.5 |
| 27 | 24 ± 1 |
| 28 | 27.7 ± 0.9 |
| 29 | 27.02 ± 0.07 |
| 30 | 25.2 ± 0.7 |
| 31 | 28.25 ± 0.08 |
| 32 | 26.7 ± 0.6 |
| 33 | 24 ± 1 |
| 34 | 26 ± 1 |
| 35 | 32 ± 1 |
| 36 | 25.2 ± 0.8 |
| 37 | 26.9 ± 0.6 |
| 38 | 33.7 ± 0.7 |
| 39 | 25.7 ± 0.6 |
| 40 | 29.6 ± 0.7 |
| 41 | 24 ± 1 |
| 42 | 24 ± 1 |
| 43 | 25 ± 1 |
| 44 | 27.9 ± 0.7 |
| 45 | 25.4 ± 0.9 |
| 46 | 26 ± 1 |

| | |
|---|---|
| 47 | 31.8 ± 0.4 |
| 48 | 31 ± 2 |
| 49 | 25 ± 1 |
| 50 | 28.2 ± 0.5 |
| 51 | 32.7 ± 0.2 |
| 52 | 32.1 ± 0.2 |
| 53 | 24 ± 1 |
| 54 | 33 ± 1 |
| 55 | 23 ± 2 |
| 56 | 24 ± 2 |
| 57 | 29.7 ± 0.6 |
| 58 | 26 ± 2 |
| 59 | 25.9 ± 0.5 |
| 60 | 22 ± 1 |
| 61 | 26.9 ± 0.6 |
| 62 | 29.4 ± 0.3 |
| 63 | 30.3 ± 0.4 |
| 64 | 35.0 ± 0.8 |
| 65 | 25.4 ± 0.7 |
| 66 | 22 ± 2 |
| 67 | 26 ± 1 |
| 68 | 25.2 ± 0.6 |
| 69 | 29.6 ± 0.8 |
| 70 | 27.7 ± 0.5 |
| 71 | 27 ± 1 |
| 72 | 23 ± 1 |
| 73 | 26.2 ± 0.4 |
| 74 | 23 ± 1 |
| 75 | 28 ± 2 |
| **AVERAGE** | **26.8 ± 0.3** |
| **P-Value** | **<0.001** |

**Table S4.** Melting temperatures for 0.25 μM FRET-labeled barnase in Eco80 and HeLa80 extracted from fitting D-aA vs. temperature

| $T_M$ (°C) | | |
|---|---|---|
| | **Eco80** | **HeLa80** |
| **Trial 1** | 52 ± 1 | 39.0 ± 0.6 |
| **Trial 2** | 49.2 ± 0.8 | 41.3 ± 0.5 |
| **Trial 3** | 52 ± 1 | 40.3 ± 0.5 |
| **Trial 4** | | 39.9 ± 0.4 |
| **Average** | 51.0 | 40.1 |
| **Standard Error** | 0.9 | 0.5 |
| **P-Value** | 0.004 | 0.001 |

**Table S5.** Melting temperatures for 0.25 μM FRET-labeled barnase in 20 mM sodium phosphate, pH 7.4, with different buffer additives extracted from fitting D-aA vs. temperature

| $T_M$ (°C) | | | | | | | |
|---|---|---|---|---|---|---|---|
| | **Phosphate Buffer** | **Ficoll** | **KCl** | **M-PER** | **Salmon Sperm DNA** | **Cytoplasmic Lysate** | **Nuclear Lysate** |
| **Trial 1** | 49 ± 2 | 48 ± 1 | 51 ± 2 | 44 ± 2 | 42 ± 2 | 47 ± 1 | 44 ± 1 |
| **Trial 2** | 45 ± 2 | 49 ± 1 | 47 ± 2 | 44 ± 2 | 43.5 + 0.6 | 44.6 ± 0.6 | 41 ± 1 |
| **Trial 3** | 47 ± 1 | 49 ± 1 | 46 ± 2 | 44 ± 2 | 43 ± 1 | 50.7 ± 0.8 | 40 ± 1 |
| **Trial 4** | 42 ± 2 | | 48 ± 1 | | | | |
| **Trial 5** | 43 ± 2 | | 50 ± 1 | | | | |
| **Trial 6** | 44 ± 2 | | 49 ± 1 | | | | |
| **Trial 7** | 47 ± 1 | | | | | | |
| **Average** | 45.2 | 48.8 | 48.4 | 44.31 | 42.7 | 48 | 41.8 |
| **Standard Error** | 0.9 | 0.3 | 0.8 | 0.06 | 0.5 | 2 | 0.9 |
| **P-Value** | | 0.008 | 0.025 | 0.393 | 0.049 | 0.31 | 0.042 |

**Table S6.** Summary of melting temperatures for barnase Y13G H102A in BL21 DE3 *E. coli* extracted from fitting D/A vs. temperature

| $T_M$ (°C) | | | |
|---|---|---|---|
| | **Batch 1** | **Batch 2** | **Batch 3** |
| **Trial 1** | 25 ± 2 | 21 ± 1 | 20.7 ± 0.4 |
| **Trial 2** | 24.8 ± 0.1 | 25 ± 2 | 21.0 ± 0.7 |
| **Trial 3** | 25.9 ± 0.6 | 26 ± 2 | 23.4 ± 0.5 |
| **Batch Average** | 25.4 ± 0.3 | 24 ± 1 | 21.7 ± 0.8 |
| **Overall Average** | 23.7 ± 0.7 | | |

**Table S7.** Summary of melting temperatures for barnase H102A in BL21 DE3 *E. coli* extracted from fitting D/A vs. temperature

| $T_M$ (°C) | | | | |
|---|---|---|---|---|
| | **Batch 1** | **Batch 2** | **Batch 3** | **Batch 4** |
| **Trial 1** | 41.0 ± 0.3 | 45.4 ± 0.3 | 44.7 ± 0.3 | 42.0 ± 0.3 |
| **Trial 2** | 38.7 ± 0.5 | 44 ± 3 | 43.2 ± 0.4 | 41.4 ± 0.3 |
| **Trial 3** | 39.2 ± 0.5 | 51 ± 1 | 41.8 ± 0.3 | 40.8 ± 0.4 |
| **Batch Average** | 39.6 ± 0.7 | 47 ± 2 | 43.2 ± 0.8 | 41.4 ± 0.4 |
| **Overall Average** | 42.7 ± 0.9 | | | |
